# Translational control of Bcl-2 promotes apoptosis of gastric carcinoma cells

**DOI:** 10.1101/608836

**Authors:** Shuangfen Tao, Jianchun Gu, Qing Wang, Leizhen Zheng

## Abstract

Aberrantly expressed microRNAs (miRNAs) are essential for the tumorigenesis of gastric carcinoma (GC). Nevertheless, an effect of miR-383 on GC cells apoptosis has not been acknowledged previously. Here, we investigated the miR-383 level and anti-apoptotic protein Bcl-2 level in specimens of GC. Bioinformatics analyses was performed, and assay of luciferase-reporter was used for analyzing the relationship between Bcl-2 and miR-383. We analyzed survival of GC cells upon Fluorouracil treatment in an assay of CCK, and measured apoptosis of cells using flow cytometric FITC Annexin V apoptosis detection assay. The level of miR-383 was found extremely lower while the level of Bcl-2 levels was found extremely higher in GC specimens, in comparison with tissue from the adjacent non-tumor region. Additionally, the miR-383 level and Bcl-2 level inversely correlated in specimens of GC. In comparison with miR-383-high subjects, MiR-383-low subjects showed overall lower survival. Bioinformatics analyses revealed that miR-383 targeted the 3’-UTR of Bcl-2 mRNA to restrain its translation, demonstrated in a luciferase reporter assay. MiR-383 overexpression inhibited the Bcl-2-mediated survival of cell against apoptosis induced by Fluorouracil, while miR-383 depletion enhanced the cell survival. Together, these data indicate that suppression of miR-383 in GC improves the Bcl-2-mediated cell survival of GC against the chemotherapy-induced cell death. MiR-383 re-expression in cells of GC might augment apoptosis of GC cells during chemotherapy.

## Introduction

In China, gastric carcinoma (GC) is the leading cause of cancer-related death [1]. Chemotherapy, such as Fluorouracil (5-FU) treatment, which has been used to surgically remove primary cancer as a supplementary treatment, substantially improves patients’ survival [2]. Of note, we have shown resistance of some GCs to 5-FU treatment while the molecular mechanisms underlying this phenomenon remains elusive. The resistance of a particular cancer to a particular chemotherapy might increase anti-apoptotic potentials of the cancer cells in special circumstances [3–5]. Apoptosis of a cell is regulated by proteins associated with control of apoptosis such as Bid, Bak, Bad, and Bcl-2 [6].

Researchers have made a lot of efforts to decipher the molecular carcinogenesis of GC, in order to find an effective therapy of molecular for assisting the surgery as well as chemotherapy [7]. Strong evidence supports contribution of abnormal MicroRNAs (miRNAs) to progression, initiation, metastases, outgrowth and resistance of GC to chemotherapy [8–10]. MiRNA is a group of small non-coding RNA of comprising of 18-23 nucleotides, and regulate the expression of gene at the level of protein, via pairing with the 31-untranslated region (31-UTR) of the target gene mRNA [11, 12]. Levels of protein are controlled by protein translation and protein degradation, in addition to gene expression. Therefore, miRNAs are crucial for many biological events that regulate protein levels including tumorigenesis [8–10, 13]. Interestingly, miR-383 has been shown to inhibit retinal pigment epithelial cell viability and promote apoptosis and ROS formation likely through Bcl-2 and peroxiredoxin 3 (PRDX3) [14]. In another study, miR-383 upregulation was shown to prevent propofol-induced apoptosis of hippocampal neuron and cognitive impairment via Bcl-2 [15]. Nevertheless, the involvement of miR-383/Bcl-2 in the anti-apoptotic carcinogenesis in GC was not reported before and was studied here.

## Materials and Methods

### Specimen of patient tissues

In this research, we used clinically and histologically diagnosed 50 resected specimens of GC (paired GC and the adjacent non-tumor gastric tissue (NT)) at the Xin Hua Hospital from 2011 to 2014. The access and application of these clinical materials for research obtained prior consent from patient and authorization from the Institutional Research Ethics Committee.

### Reagents and cell line

We purchased 2 human GC cell lines, including AGS and SNU-5, from American Type Culture Collection (ATCC, Rockville, MD, USA). We cultured these two cell lines in RPMI1640 medium (Invitrogen, Carlsbad, CA, USA) with 10% fetal bovine serum supplementation (FBS; Sigma-Aldrich, St Louis, MO, USA) in 5% CO_2_ humidified chambers under temperature 37 °C. Fluorouracil (5-FU; Sigma-Aldrich) was diluted in 1mmol/l stock and 5µmol/l was applied to GC cells in culture.

### Plasmid transfection

We purchased plasmids carrying antisense (as) for miR-383 and miR-383-expressing (RiboBio Co., Ltd., Guangzhou, Guangdong, China), using in a transfection employing Lipofectamine 3000 reagent (Invitrogen), following manufacturer’s protocol. We transfected the cells with null plasmid as a control (null) and purified either transfected cells expressing miR-383, or as-miR-383, or control null by flow cytometric analysis for GFP.

### ELISA

We extracted protein from the specimens of both GC or NT, or cultured cells, in RIPA lysis buffer (0.1% SDS, 1% NP40, 0.5% sodium deoxycholate, 100μg/ml, phenylmethylsulfonyl fluoride, in PBS) on ice. The ELISA for Bcl-2 was performed using a human Bcl-2 ELISA kit (LS-F4134, LSbio Lifespan Biosciences Inc, Seattle, WA, USA).

### Quantitative RT-PCR (RT-qPCR)

We extracted total RNAs from cultured cells or tissue specimens using miRNeasy mini kit (Qiagen, Hilden, Germany). Complementary DNA (cDNA) was primed randomly with High-Capacity cDNA Reverse Transcription Kit (Applied Biosystems, Foster City, CA, USA) initiating with 2μg of total RNA. RT-qPCR was performed subsequently using a QuantiTect SYBR Green PCR Kit (Qiagen). We purchased all primers from Qiagen, collected raw data and analyzed levels of relative mRNA expression with 2-ΔΔCt method. Gene values were obtained by sequential normalization against housekeeping gene GAPDH and the control of the experiments.

### Prediction of MiRNA target and assay of 3’-UTR luciferase-reporter

As mentioned before, MiRNAs targets were predicted with the algorithms TargetSan (https://www.targetscan.org) [16]. Molecular cloning technology was successfully used to construct Luciferase-reporters. We purchased Target sequences for Bcl-2 miRNA 3’-UTR clone and the one with a site mutation at miR-383-binding site from Creative Biogene (Shirley, NY, USA). We seeded MiR-383-modified cells of GC in 24-well plates, and then transfected the cells with 1μg of Luciferase-reporter plasmids each well after 24 hours. Dual-luciferase reporter gene assay kit (Promega, Beijing, China) was used to measure activities of Luciferase, following the instructions of the manufacturer.

### Cell counting kit-8 (CCK-8) assay

Cell viability was measured using the CCK-8 detection kit (Sigma-Aldrich) according to the protocol of manufacturer.

### The assay of apoptosis by flow cytometry

We re-suspended the cultured cells or dissociated tissue cells in PBS to analyze cell proliferation. Cells underwent FAC analysis to determine the percentage of Annexin V+ PI-apoptotic cells after double staining using FITC-Annexin V and propidium iodide (PI) from a FITC Annexin V Apoptosis Detection Kit I (Becton-Dickinson Biosciences, San Jose, CA, USA).

### Analysis of Statistics

All of the data in this study were analyzed statistically with a one-way ANOVA method coupled with a Bonferroni correction. Fisher’s Exact Test was used for comparison between two groups (GraphPad Prism 7, GraphPad Software, Inc. La Jolla, CA, USA). We used Spearman’s Rank Correlation Coefficients to calculate Bivariate correlations. We used Kaplan-Meier curves to analyze survival of patient grouped according to the levels of miR-383. All values were expressed as mean ± standard deviation and considered significant if p < 0.05.

## Results

### Alteration in miR-383 and Bcl-2 inversely correlates in specimens of GC

We analyzed 50 specimens of GC and found that Bcl-2 levels in GC specimens were significantly higher (Figure 1A-B), while miR-383 levels in GC specimens were significantly lower (Figure 1C-D), comparing with the tissues of adjacent non-tumor gastric (NT) from the same patient. In order to examine the relationship between Bcl-2 and miR-383, we investigated the levels of both miR-383 and Bcl-2 in the 50 GC specimens, from which we detected a strong inverse correlation between the two factors (Figure 1E, 1= −0.76, p<0.0001). Our data thus revealed a causal link between miR-383 and Bcl-2 in GC cells. Moreover, we examined if the miR-383 levels might correlate with the overall GC patients’ survival. We followed the fifty patients up for five years and selected the median value of all fifty cases as the cutoff point to separate the miR-383-high cases (n=25) from the miR-383-low cases (n=25). Kaplan-Meier curve was performed to show that the overall survival of miR-383-low GC patients was obviously poorer than miR-383-high GC patients (Figure 1F).

**Figure 1:**
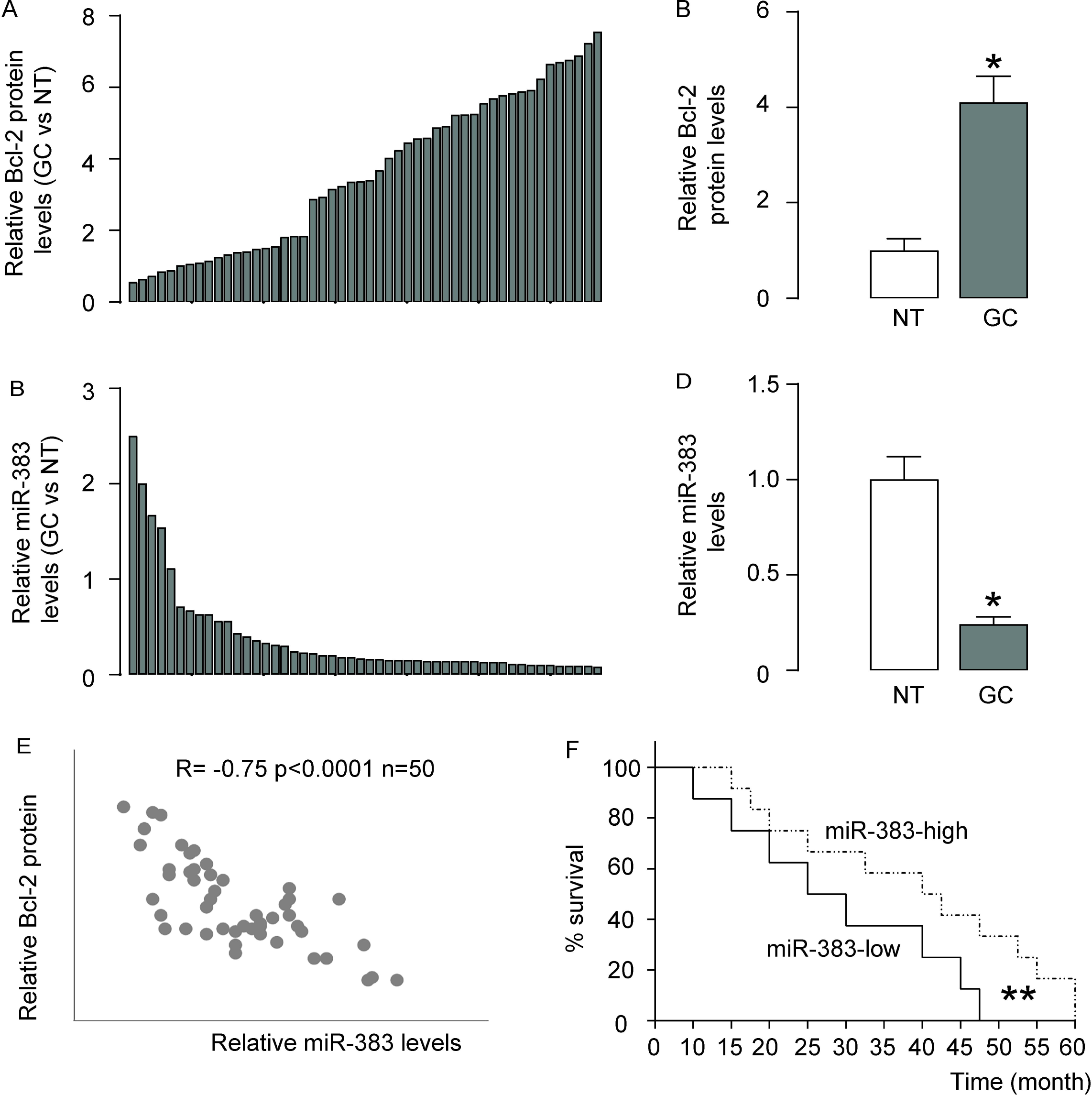
Decreased miR-383 and correlated increased Bcl-2 are detected in GC. RT-qPCR on miR-383 and ELISA for Bcl-2 were performed on paired GC and the adjacent non-tumor gastric tissues (NT) from 50 patients. (A-B) Bcl-2 levels, shown by individual values (A) and by summary (B). (C-D) miR-383 levels, shown by individual values (C) and by summary (D). (E) A Correlation test between Bcl-2 and miR-383. (F) The 50 patients were followed-up for 5 years. The median value of all 50 cases was chosen as the cutoff point for separating miR-383-high cases (n=25) from miR-383-low cases (n=25). Kaplan-Meier curves were performed to evaluate the overall survival of the GC patients, based on miR-383 levels. *p<0.05. N=50.

### MiR-383 suppresses translation of Bcl-2 via targeting 3’-UTR mRNA

A relationship between Bcl-2 and miR-383 in cells of GC was suggested from above data, which inspired us to examined whether miR-383 may target Bcl-2 mRNA to inhibit its translation. A miR-383-binding site at the 3’-UTR of Bcl-2 mRNA ranged from 427 to 433 base site was predicted by analyses of Bioinformatics (Figure 2A). We either overexpressed or inhibited miR-383 in AGS and SNU-5 human GC cell lines, by transfecting the cells with plasmids carrying either a miR-383-mimic (AGS-miR-383; SNU-5-miR-383), or a miR-383 antisense (AGS-as-miR-383; SNU-5-as-miR-383), to figure out if the binding of miR-383 to Bcl-2 mRNA in cells of GC could be functional. Also, the AGS and SNU-5 cells were transfected with a plasmid carrying null sequence as a control (AGS-null; SNU-5-null). RT-qPCR confirmed the overexpression and inhibition of miR-383 in the AGS and SNU-5 cells (Figure 2B-C). Subsequently, we transfected miR-383-modified cells with Bcl-2 3’-UTR luciferase-reporter plasmid, with or without a site mutation on the miR-383 binding sites. The activities of luciferase were quantified in these cells, which suggested that the binding of miR-383 to 3’-UTR of Bcl-2 mRNA inhibited the translation of Bcl-2 protein (Figure 2D-E).

**Figure 2:**
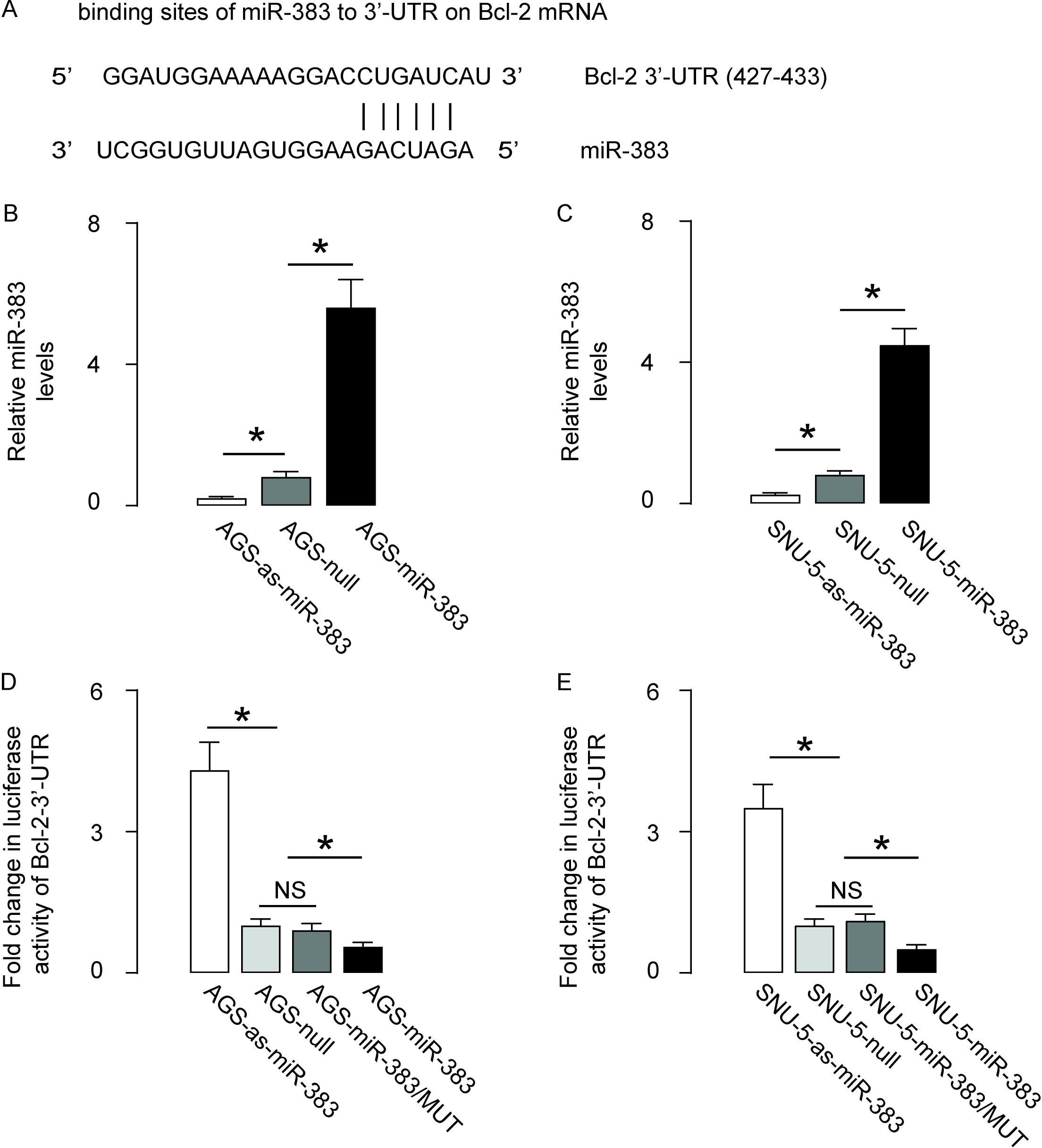
MiR-383 targets 3’-UTR of Bcl-2 mRNA to inhibit its expression. (A) Bioinformatics analyses for binding of miR-383 onto the 3’-UTR of Bcl-2 mRNA. (B) We either overexpressed miR-383, or inhibited miR-383 in a human GC cell line, AGS, by transfection of the cells with a miR-383-expressing plasmid (AGS-miR-383), or with a plasmid carrying miR-383 antisense (AGS-as-miR-383). The AGS cells were also transfected with a plasmid carrying a null sequence as a control (AGS-null). The modification of miR-383 levels in AGS cells was confirmed by RT-qPCR. (C) We either overexpressed miR-383, or inhibited miR-383 in a human GC cell line, SNU-5, by transfection of the cells with a miR-383-expressing plasmid (SNU-5-miR-383), or with a plasmid carrying miR-383 antisense (SNU-5-as-miR-383). The AGS cells were also transfected with a plasmid carrying a null sequence as a control (SNU-5-null). The modification of miR-383 levels in SNU-5 cells was confirmed by RT-qPCR. (D) MiR-383-modified AGS cells were transfected with 1μg of Bcl-2-3’UTR luciferase-reporter plasmid. The luciferase activities were examined. (E) MiR-383-modified SNU-5 cells were transfected with 1μg of Bcl-2-3’UTR luciferase-reporter plasmid. The luciferase activities were examined. *p<0.05. N=5.

### MiR-383 decreases Bcl-2 protein in cells of GC with affecting mRNA

We found that miR-383 level modification in either AGS cells (Figure 3A) or SNU-5 cells (Figure 3B) failed to alter Bcl-2 mRNA levels. Nevertheless, miR-383 overexpression significantly decreased levels of Bcl-2 protein, while miR-383 depletion significantly increased levels of Bcl-2 protein in both cell lines, by ELISA (Figure 3C-D). Hence, miR-383 inhibits Bcl-2 mRNA translation in cells of GC.

**Figure 3:**
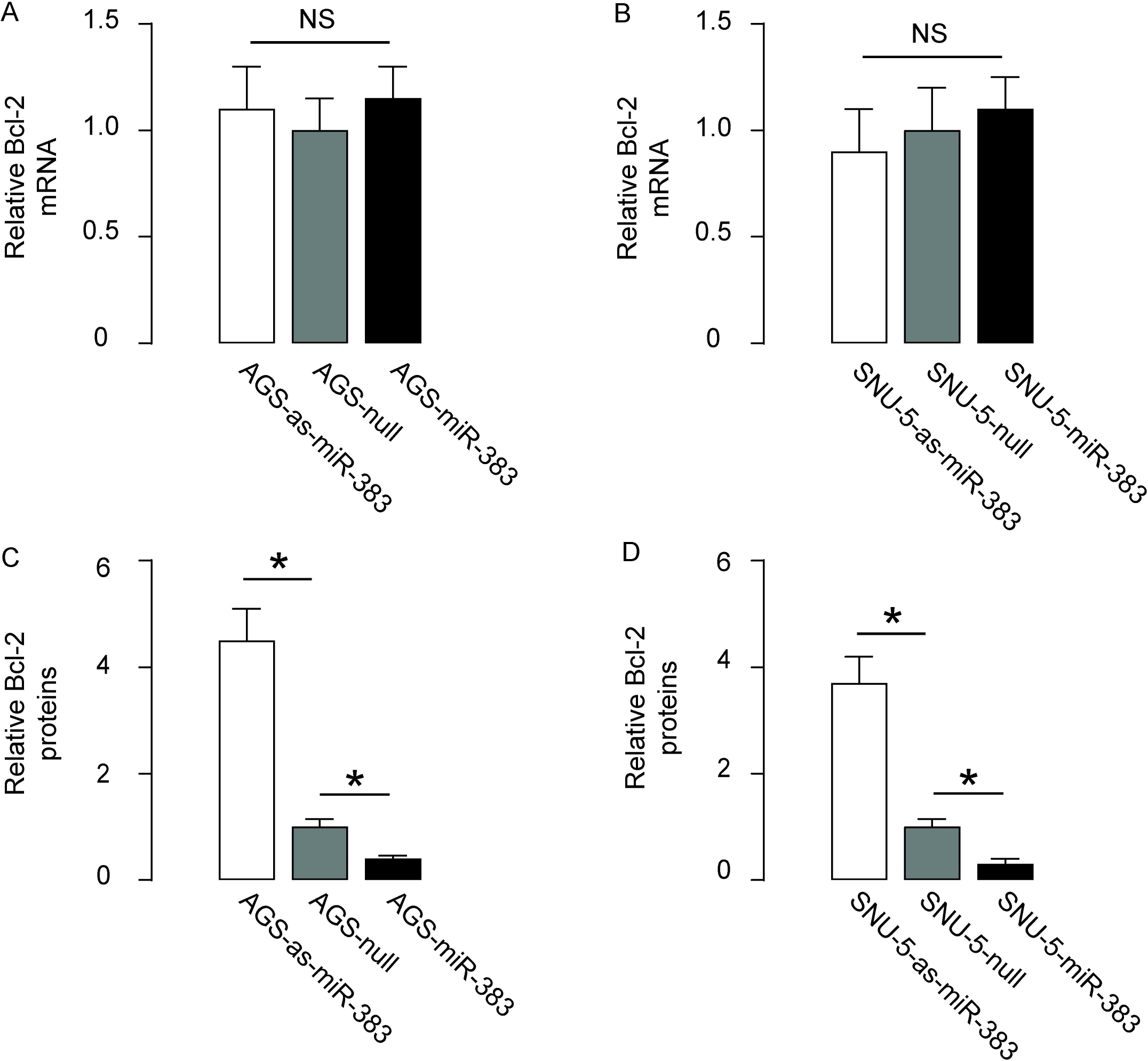
MiR-383 decreases Bcl-2 protein, but not mRNA, in GC cells. (A-B) The Bcl-2 mRNA levels in miR-383-modified AGS cells (A) and miR-383-modified SNU-5 cells (B). (C-D) The Bcl-2 protein levels in miR-383-modified AGS cells (C) and miR-383-modified SNU-5 cells (D). *p<0.05. NS: non-significant. N=5.

### MiR-383 increases 5-FU-induced apoptosis of GC cells

Finally, we investigated the effects of miR-383 modification on cell viability of GC in 5-FU challenge in an assay of CCK-8. We found that miR-383 overexpression decreased cell viability of SNU-5 and AGS cells, which was abolished by overexpression of Bcl-2 (Figure 4A-B). Moreover, miR-383 depletion increased cell viability of SNU-5 and AGS cells, which was abolished by depletion of Bcl-2 (Figure 4A-B). Next, we did assay of cytometric analysis of apoptosis, and found that miR-383 overexpression increased cell apoptosis in both AGS (Figure 4C-D) and SNU-5 (Figure 4E-F) cells, which could be abolished by overexpression of Bcl-2 (Figure 4C-F). On the other hand, miR-383 depletion decreased cell apoptosis in both AGS (Figure 4C-D) and SNU-5 (Figure 4E-F) cells, which could be abolished by depletion of Bcl-2 (Figure 4C-F). Thus, miR-383 increases 5-FU-induced apoptosis of GC cell through Bcl-2. In summary, these data indicate that miR-383 inhibition in GC promotes Bcl-2-mediated cell survival against apoptotic death induced by chemotherapy. The level of miR-383 re-expression in cells of GC could promote cancer cell apoptosis in chemotherapy (Figure 5).

**Figure 4:**
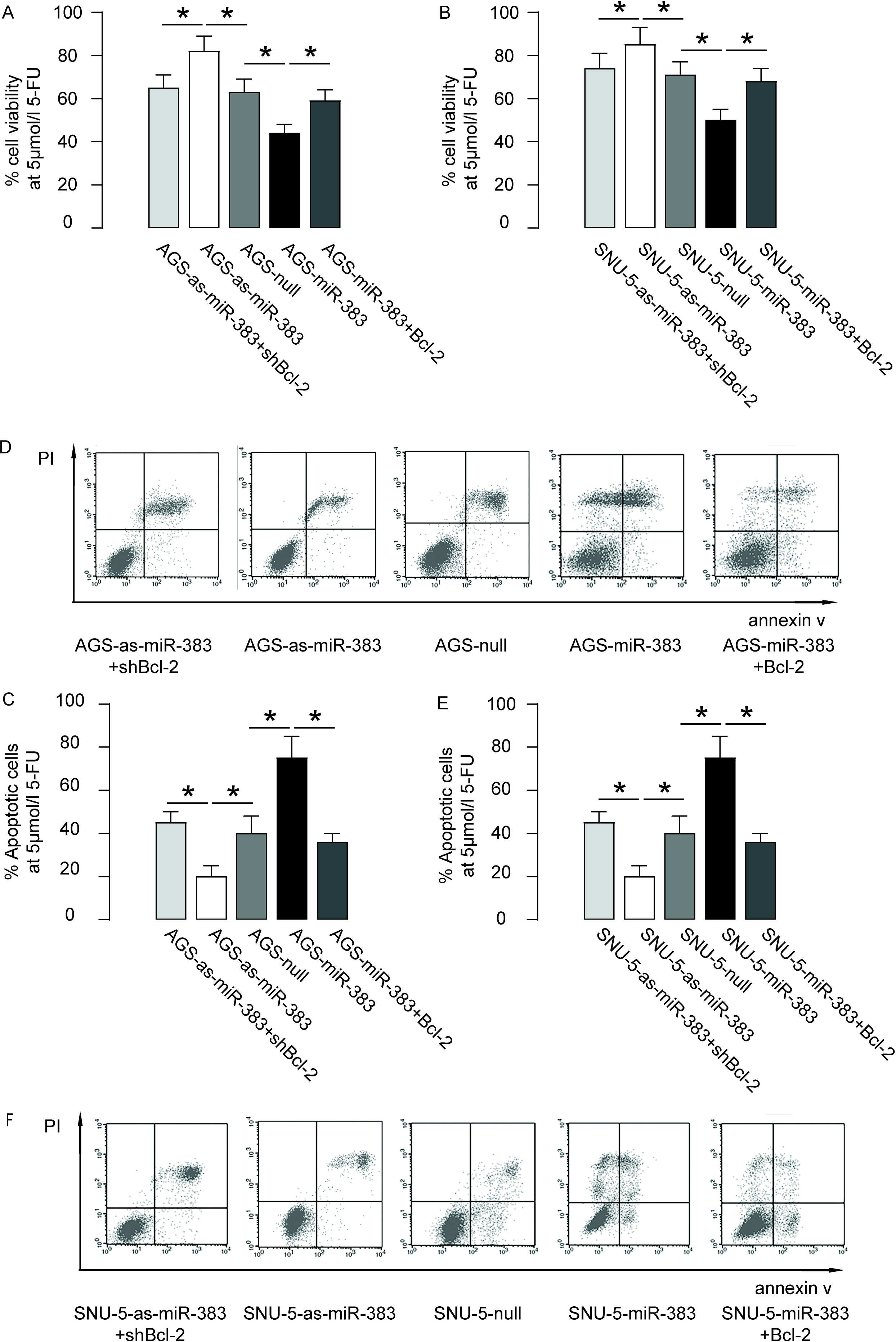
MiR-383 increases 5-FU-induced GC cell apoptosis. (A-B) The effects of miR-383 modification on AGS (A) and SNU-5 (B) cell viability at presence of 5µmol/l 5-FU in an CCK-8 assay. (C-D) Fluorescence-based apoptosis assay on miR-383-modified AGS cells, shown by quantification (C), and by representative flow charts (D). (E-F) Fluorescence-based apoptosis assay on miR-383-modified SNU-5 cells, shown by quantification (E), and by representative flow charts (F). *p<0.05. N=5.

**Figure 5:**
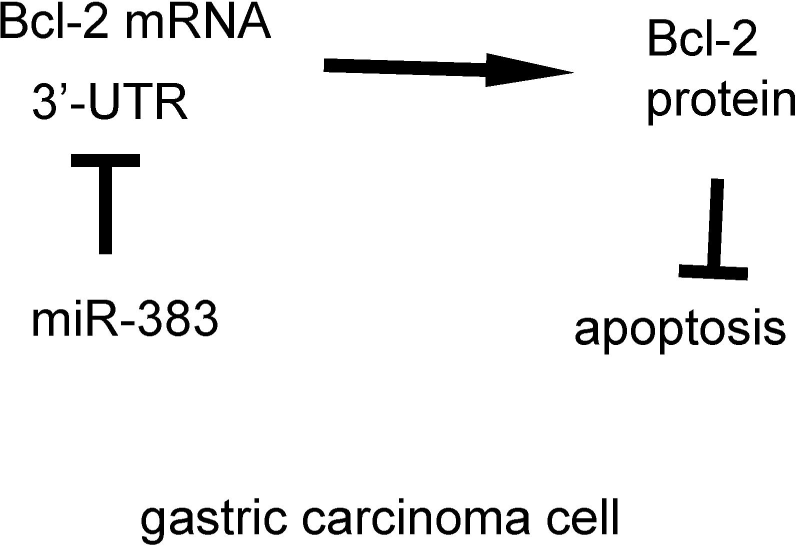
Schematic of the model. MiR-383 suppression in GC promotes Bcl-2-mediated cancer cell survival against chemotherapy-induced cell death. Re-expression of miR-383 levels in GC cells may enhance cancer apoptosis during chemotherapy.

## Discussion

Many miRNAs play an important role in different stages of GC, including initiation, invasion, metastases, progression, and resistance of chemotherapy. The realization of miRNA function could be mediated through either itself or through synergizing with other key regulators to coordinate the GC pathogenesis. Therefore, understanding of the role of aberrantly expressed miRNAs in carcinogenesis of GC might could contribute to decipher the mechanisms underlying progression of GC.

The GC cell resistance to chemotherapy is mainly due to the compromised activation of apoptotic machinery and the aberrant activation of anti-apoptotic machinery inside the cells of GC. We looked into earlier research and found that Bcl-2 activation is possibly a key reaction factor in cells of GC in 5-FU treatment, including activation of apoptosis-association proteins, such as Bak, Bid, Bad, augmentation of caspases and Cytochrome C to initiate the process. Bcl-2 inhibits these pro-apoptotic proteins efficiently, but the Bcl-2 activation in response to chemotherapy appears case-dependent.

Here, we found some candidate miRNAs that target Bcl-2 by sequence matching in bioinformatics analyses, and we detected a major decrease specifically in miR-383 in specimens of GC among these miRNAs, comparing with the tissue of paired non-tumor. Interestingly, miR-383 has been shown to inhibit retinal pigment epithelial cell viability and promote apoptosis and ROS formation likely through Bcl-2 and peroxiredoxin 3 (PRDX3) [14]. In another study, miR-383 upregulation was shown to protect hippocampal neuron from undergoing apoptosis in response to propofol via Bcl-2 [15]. Very recently, cell-cycle-related and expression-elevated protein in tumor (CREPT) was found to transcriptionally enhance Cyclin D1 expression to promote colorectal carcinoma cell replication likely by miR-383 [17]. Based on these studies, we hypothesized that miR-383 might not only target but also regulate Bcl-2 in cells of GC. This hypothesis was further supported by correlation test and by the luciferase reporter assay. Then, we modified the levels of miR-383 in cells of GC and found that it failed to affect Bcl-2 mRNA but altered protein abundancy. In addition, it seemed that miR-383-mediated changes in Bcl-2 altered the cell viability in response to 5-FU. In this report, the effects of miR-383 on cell invasion were not analyzed.

In conclusion, we presented a model to explain the mechanisms that underlie the resistance of GC cells to chemotherapy. The downregulation of miR-383 by GC cells results in strengthened translation of Bcl-2 protein translation, which subsequently enables GC cells to survive upon chemotherapy. On the other hand, re-expression of miR-383 in cells of GC might substantialize GC cell apoptosis strengthen during chemotherapy.

